# Determinants of multiheme cytochrome extracellular electron transfer uncovered by systematic peptide insertion

**DOI:** 10.1101/2022.03.09.483668

**Authors:** Ian J. Campbell, Joshua T. Atkinson, Matthew D. Carpenter, Dru Myerscough, Lin Su, Caroline M. Ajo-Franklin, Jonathan J. Silberg

**Author notes:** Jonathan J. Silberg, Department of Biosciences, Rice University, 6100 Main Street, MS-140, Houston, TX, 77005. **Author Contributions:** I.J.C., J.T.A., and J.J.S. conceptualized the project. I.J.C. and J.T.A. constructed the library. I.J.C. performed selections. I.J.C. and M.D.C. performed ECL analysis. I.J.C., L.S., and M.D.C. performed nanoparticle assays. M.D.C. conducted the electrochemistry measurements. I.J.C. and D.M. conducted structural analysis. I.J.C. and J.J.S. drafted the article and all authors contributed text.

## Abstract

The multiheme cytochrome MtrA enables microbial respiration by transferring electrons across the outer membrane to extracellular electron acceptors. While structural studies have identified residues that mediate MtrA binding to hemes and to other cytochromes that facilitate extracellular electron transfer (EET), the relative importance of these interactions for EET is not known. To better understand EET, we evaluated how insertion of an octapeptide across all MtrA backbone locations affects *Shewanella oneidensis* MR-1 respiration on Fe(III). EET efficiency was found to be inversely correlated with insertion proximity to the heme prosthetic groups. Mutants with decreased EET also arose from insertions in a subset of the regions that make residue-residue contacts with the porin MtrB, while all sites contacting the extracellular MtrC presented high peptide insertion tolerance. MtrA variants having peptide insertions within the CXXCH motifs that coordinate heme cofactors retained some ability to support respiration on Fe(III), although these variants presented significantly decreased EET. Furthermore, the fitness of cells expressing different MtrA variants under Fe(III)-respiring conditions correlated with anode reduction. The peptide-insertion profile, which represents the first comprehensive sequence-structure-function map for a multiheme cytochrome, implicates MtrA as a strategic protein engineering target for regulating EET.

## INTRODUCTION

Multiheme cytochrome electron transfer proteins have evolved to support the exchange of electrons between cells and the extracellular environment.^1, 2^ By binding a series of adjacent cofactors that span tertiary structures, these proteins enable electron transport across the insulating membrane and across micron-scale distances by insulating the heme prosthetic groups from the solvent.^3–6^ This high efficiency electron transport enables diverse microorganisms to respire on extracellular metals, minerals, and materials.^2, 7–9^ The Gram-negative bacterium *Shewanella oneidensis* has evolved a cytochrome-based pathway, designated the metal-reducing pathway (Mtr) to support extracellular electron transfer (EET).^10, 11^ This pathway allows microbes to respire on Fe(III), Mn(III), and Mn(IV).^10, 11^ Additionally, the Mtr pathway can reduce extracellular minerals, like hematite or ferrihydrite, as well as synthetic nanoparticles and electrodes.^7, 12–17^

Among the different EET systems that have been reported, the Mtr pathway of *Shewanella oneidensis* has been studied most intensively.^18–20^ In this pathway, multiheme cytochromes housed within porins transfer reducing equivalents generated by catabolic processes through the outer membrane.^21–23^ While *S. oneidensis* encodes multiple cytochrome-porin complexes, the complex with the highest flux of electrons is MtrCAB (UniProt Q8EG34, Q8EG35, and Q8CVD4).^22^ MtrC is required for appreciable reduction of ferric citrate, iron oxides, and flavins.^15, 22^ The loss of MtrC leads to a large loss in current supplied to an electrode and decreased reduction of synthetic materials, including UO_2_ and Pd nanoparticles.^15–17^ Similarly, loss of MtrA decreases the ability of *S. oneidensis* to reduce ferric citrate, iron oxide, and flavins, an effect that is amplified when coupled with knockouts of MtrA paralogs.^22^

Structural studies have provided atomistic insight into molecular interactions within the MtrCAB complex.^20, 24^ The decaheme cytochrome MtrA resides within MtrB, a porin that spans the extracellular membrane. The decaheme MtrC binds to the extracellular face of the MtrAB complex and supports electron transfer to exogenous electron acceptors by offering a large surface area with multiple surface exposed hemes.^20, 22^ Spectroscopic and simulation studies have found that electrons can pass between adjacent heme prosthetic groups within the MtrCAB complex at nanosecond rates and suggested that these rapid transfers are facilitated by cysteine linkages positioned between those tetrapyrroles.^3, 4, 6^ Interestingly, a cysteine linkage has also been observed between heme A10 of MtrA and heme C5 of MtrC; electron transfer between these hemes is thought to be rate limiting for EET through the MtrCAB complex.^4^ While these studies have revealed a complex network of molecular interactions within the Mtr complex, including protein-cofactor and protein-protein interactions, we cannot yet anticipate how changes to the primary structure of MtrCAB affect EET.

One way to probe the importance of native residues in mediating protein-cofactor and protein-protein interactions is to evaluate the effects of mutations on cellular function.^25–27^ In cases where a high-throughput selection is available, large numbers of mutations can be evaluated in parallel using laboratory evolution.^26–28^ With this approach, a library of genes encoding mutant proteins is created, and the library is deep sequenced before and after selection for biomolecules with parent-like functions.^29^ The changes in the abundance of each mutant are then used to calculate the ratio of sequence counts before and after selection, which can be used to estimate the relative activities of mutant proteins. This approach has been applied to diverse proteins that function in cellular catalysis.^30–33^ However, it has not yet been applied to multiheme cytochromes that mediate EET.

To study sequence-structure-EET relationships in MtrA, we created a library of variants with the peptide SGRPGSLS inserted at every backbone position and selected this library for variants that support EET. We generated insertional mutations rather than amino acid substitutions because they have the potential to create a higher level of structural disruption at each native site due to the need to accommodate a large change in primary structure. By quantifying variant abundances in our library before and after selection, we identify regions that present varying EET efficiency following peptide insertion. We show that the sequence enrichments observed in our selection correlate with EET of individual variants and evaluate how peptide insertion in partner-binding regions and CXXCH motifs affect EET to Fe(III), nanoparticles, and electrodes.

## RESULTS

### MtrA library design and selection

In a prior study, a *S. oneidensis* mutant was created that lacks MtrA, three MtrA paralogs (MtrD, DmsE, SO4360), and a periplasmic electron carrier (CctA).^22^ This strain, designated *So*-*ΔmtrA* herein, is severely diminished in its ability to reduce ferric citrate. Because *S. oneidensis* is able to respire on extracellular sources of iron using the Mtr pathway, we reasoned that *So*-*ΔmtrA* growth could be complemented by MtrA expression under anaerobic conditions when ferric citrate is provided as a terminal electron acceptor.^11, 22^ To test this idea, we created plasmids that express MtrA using the *P_LtetO-1_* promoter, transformed *So*-*ΔmtrA* with these plasmids, and evaluated growth under selective (+ferric citrate) and non-selective (+fumarate) growth conditions. Under selective conditions, growth was not observed with *So*-*ΔmtrA* harboring an empty vector (Figure 1a). In contrast, robust growth was observed with *So*-*ΔmtrA* expressing MtrA, which was similar to that observed with wildtype *S. oneidensis* MR-1 expressing MtrA. All strains grew to a similar extent under non-selective conditions when provided with fumarate as an electron acceptor, which is directly reduced by the periplasmic cytochrome FccA without the need for EET through MtrA.^34^ Taken together, these findings suggest that *So*-*ΔmtrA* can be used to mine combinatorial libraries of MtrA variants for multiheme cytochromes that support EET.

**Figure 1.**
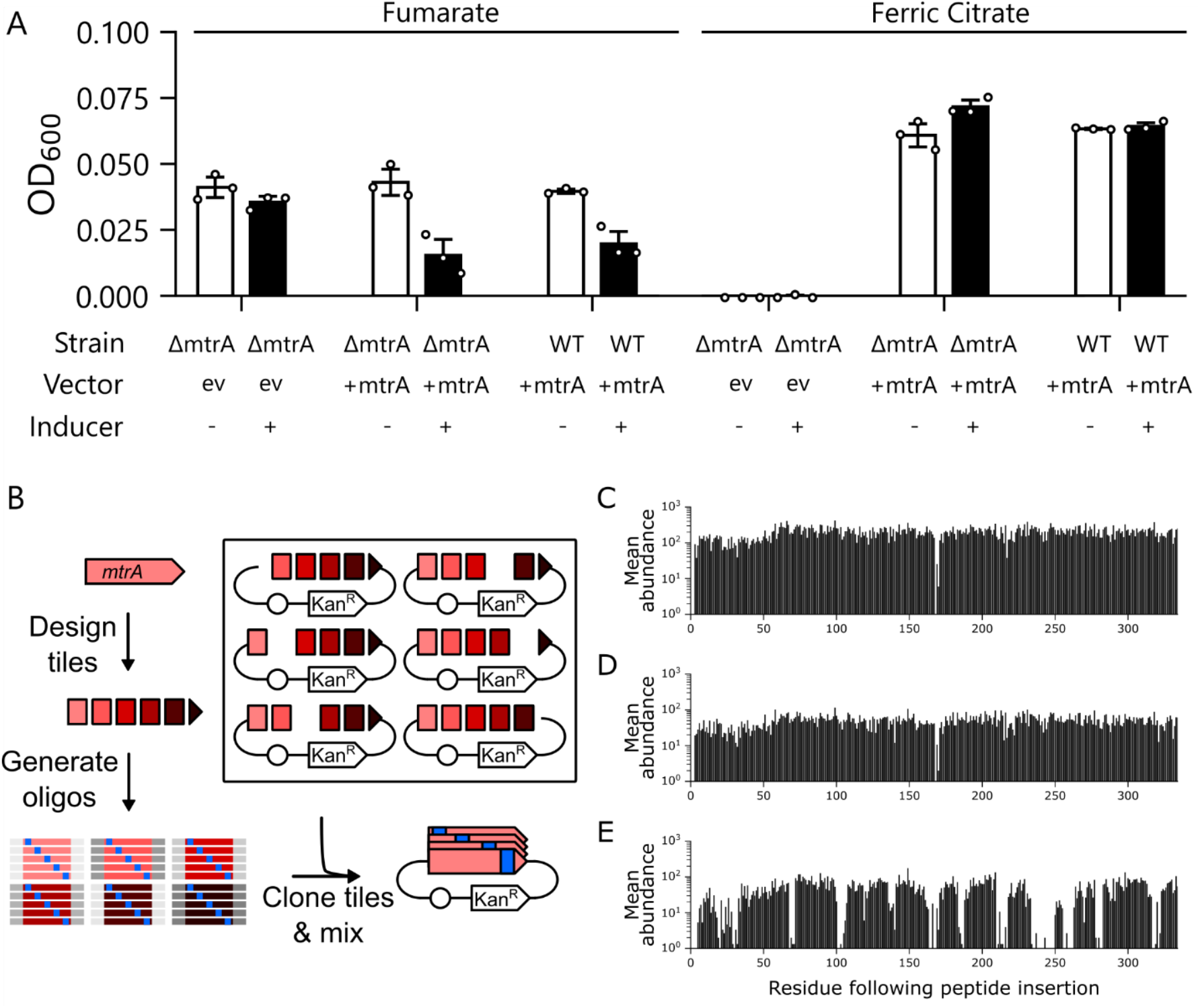
Using laboratory evolution to evaluate MtrA tolerance to peptide insertion. (**A**) *S. oneidensis So*-*ΔmtrA* requires MtrA for growth complementation on Fe(III). *S. oneidensis So*-*ΔmtrA* strains transformed with an empty vector (ev) or a plasmid expressing MtrA (+mtrA) were grown for 48 hours at 30°C in an anaerobic atmosphere with using either fumarate (white) or ferric citrate (black) as the sole electron acceptor. As a control, we also evaluated *S. oneidensis* MR-1 expressing MtrA. Error bars represent standard deviation calculated from three experiments. With ferric citrate, cells expressing MtrA grew to a significantly higher density than cells containing an empty vector (P < 0.01; two-tailed t-test), and the growth of these cells was not significantly different from *S. oneidensis* MR-1 expressing MtrA (P > 0.1; two-tailed t-test). (**B**) Using the SPINE algorithm, the MtrA sequence was fragmented into six tiles *in silico*, each having a length (<300 base pairs) that could be commercially synthesized using pooled oligonucleotide synthesis. The algorithm designed oligonucleotides encoding every unique eight codon insertion within the tile, and added unique barcodes to the ends of each tile ensemble to allow for selective amplification. PCR amplified tiles were cloned into vector backbones complementary to each tile using golden gate cloning to generate an insertion library. Deep sequencing was used to determine the average frequency (*f_i_*) of insertion sites across the MtrA library in (**C**) the naïve preculture, (**D**) cells growth anaerobically with fumarate, and (**E**) cells grown anaerobically with ferric citrate. For each condition, we show the average of three separate experiments.

To design a library of vectors that express MtrA variants with a peptide inserted at different native sites, we used a previously described algorithm called SPINE.^28^ Oligonucleotides were synthesized that encode MtrA variants with a twenty four base pair insertion, encoding SGRPGSLS, inserted between every codon. This vector library was built using a tile based cloning approach (Figure 1b). With this approach, the full length MtrA gene was broken into six tiles, and ensembles of synthetic DNA (∼230 base pairs) were cloned in parallel to replace the native sequence at different locations. To achieve a high transformation efficiency of *So*-*ΔmtrA* critical to library sampling, we demethylated the plasmid library by first transforming it into *Escherichia coli* lacking the Dam and Dcm methylation machinery.^35, 36^ The demethylated library was then electroporated into *So*-*ΔmtrA*. This protocol yielded a CFU count (∼20,000) that is predicted to sample >99% of all variants in the library.^37^ *So*-*ΔmtrA* transformed with the library were harvested, pooled, and cultured aerobically in triplicate. Aliquots of each culture (∼10^6^ CFU) were used to inoculate growth medium (100 mL) containing ferric citrate or fumarate, and these cultures were incubated under anaerobic conditions. Large cultures were used to ensure that cells would undergo a large number of doublings as they grew to stationary phase, thereby enabling the discovery of MtrA variants with a wide range of EET activities.

To examine the effect of growth complementation on MtrA mutant abundance, deep sequencing was used to quantify the relative abundance of each variant grown in: (i) the naïve aerobic *So*-*ΔmtrA* cultures used for inoculation, (ii) the non-selective anaerobic fumarate cultures, and (iii) and the selective anaerobic ferric citrate cultures. We first evaluated how the sequence biases in the inoculum compared to libraries created using transposase mediated insertion,^28, 38–41^ which can present large sequence biases in naïve libraries. Deep sequencing revealed that all of the variants were present in the naïve library except the variant having the peptide inserted before Val168 (Figures 1c, S1). The average abundance of each unique variant was 3 per 1000 sequence reads with a standard deviation of 1.2 per 1000 reads. The coefficient of variance for mutant abundance (CV = 0.37; Figure S2a) was much smaller than that observed in libraries created using transposases.^38–41^ Sequencing revealed similar relative abundances of insertion variants in samples after anaerobic growth with fumarate (Figures 1d, S3). The non-selective conditions yielded a similar CV (0.39) as the naïve library (Figure S2b). Additionally, the average abundance of each variant in the naïve and non-selective conditions exhibited a linear correlation (Figure S4). This finding suggests that MtrA mutations do not have large effects on *So*-*ΔmtrA* fitness when using fumarate as a terminal electron acceptor. Analysis of the MtrA sequence diversity in cells cultured on ferric citrate revealed more dramatic changes in the relative abundances of each variant (Figures 1e, S5), with multiple variants being absent. Under selective conditions, a skewed distribution was observed with a CV of 0.82 (Figure S2c). These findings suggest that a subset of the peptide insertions alter EET mediated by MtrA.

To compare the effects of peptide insertion at different backbone locations on Mtr-mediated EET, we calculated the relative frequency of each variant (*f_i_*) by quantifying the ratio of the reads for that variant (*r_i_*) to the total sample reads (*r_total_*) in the different growth conditions. To establish the enrichment of each variant under selective (ε_S_) and non-selective (ε_N_) conditions, we then calculated the ratio of *f_i_* obtained under selective conditions (^S^*f_i_*) to the naïve condition (^N^*f*_i_) and the ratio of *f_i_* obtained under non-selective conditions (^NS^*f_i_*) to the naïve condition (^N^*f*_i_). Z scores were then calculated for each variant by subtracting the mean ε_N_ for all variants from the ε_S_ values for the individual variants and normalizing the resulting values to the standard deviation of the ε_N_ values. The sequence enrichment varied across the primary structure. The most highly enriched sequences on ferric citrate had a z score of 7.57 (Figure 2a), while the most depleted sequences had z scores of -4.47; variants with the lowest z scores were not observed by deep sequencing following the ferric citrate selection. Mapping these z scores onto the *Shewanella baltica* MtrA crystal structure reveals that regions of low and high peptide insertion tolerance are dispersed across the tertiary structure (Figure 2b). In summary, a comparison of the sequencing data from each growth condition identified MtrA variants that complement *So*-*ΔmtrA* to differing extents, implicating these variants as transferring electrons across the outer member with a range of efficiencies.

**Figure 2.**
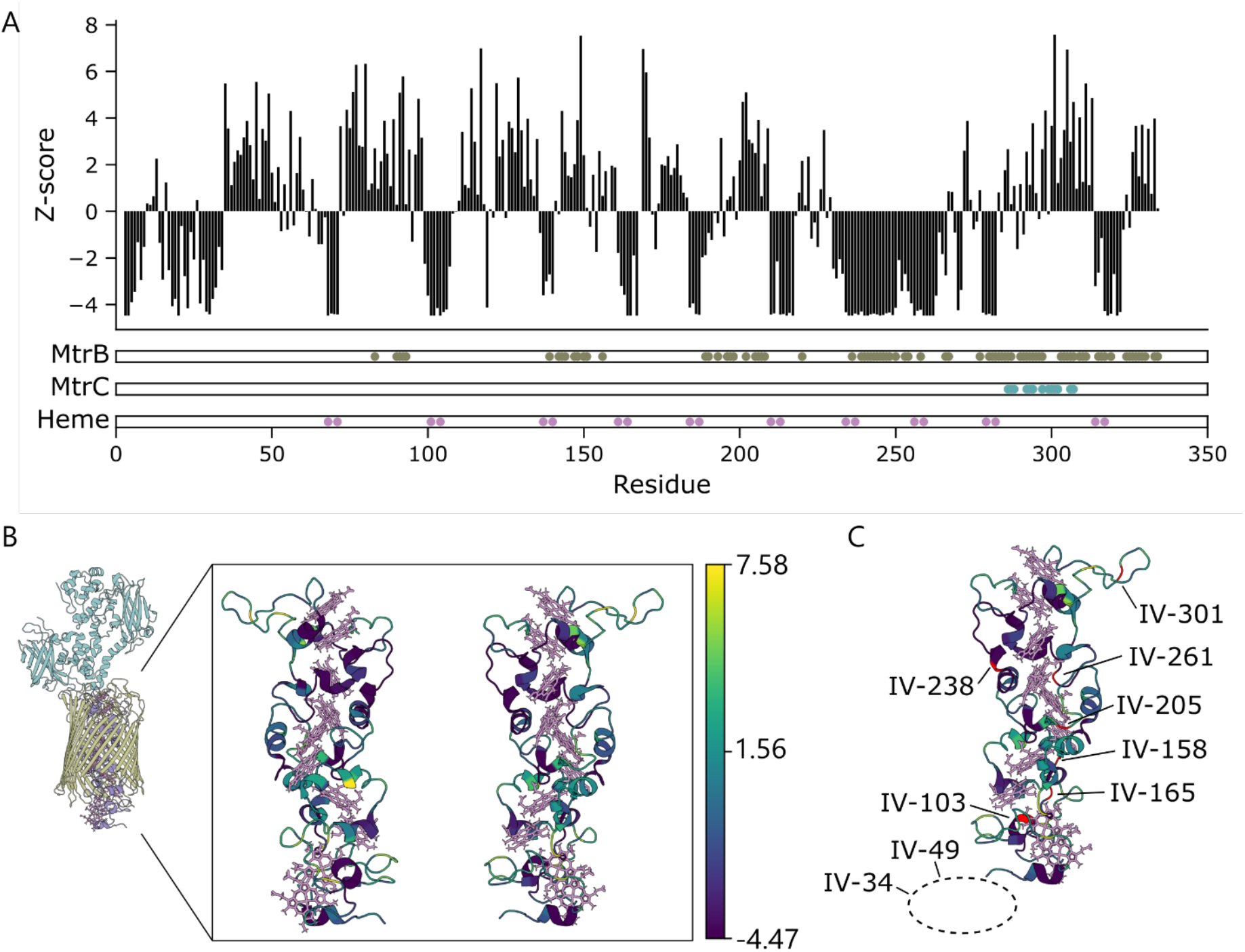
Effect of the growth selection on MtrA mutant abundances. (**A**) Enrichment values for the non-selected fumarate (^n^ε_i_) and selected ferric citrate (^s^ε_i_) cultures were calculated by dividing the frequencies observed for each mutant in the different growth conditions by the frequencies present in the naïve aerobic condition. The distribution of z-scored selected enrichments were then calculated from the selected enrichments (^s^ε_I_) normalized by the non-selective enrichment distribution (^n^ε). The MtrA residues that make contacts with MtrB (dark green) and MtrC (teal) are colored, as well as the residues that ligate hemes (pink). A distance cutoff of 8Å was used to define contacts. (**B**) The MtrA structure is color coded using the z scores observed in the mutational profile, and (**C**) the selected insertion variants of MtrA were colored red and labeled on the MtrA structure.

## MtrA mutant enrichments correlate with respiration on ferric citrate

To understand how the z scores observed for different variants relate to EET mediated by specific MtrA mutants, we chose six variants to characterize individually. For these measurements, we chose peptide insertion variants (IV) with the full spectrum of z scores (Table 1; Figures 2c, S6). These variants sample MtrA regions that make a high density of contacts with MtrB (IV-158, IV-205, IV-261) and MtrC (IV-301), as well as the structurally-unresolved N-terminal region that extends into the periplasm (IV-34, IV-49). Expression vectors for each variant were transformed into *So*-*ΔmtrA* and growth complementation was measured in anaerobic cultures containing ferric citrate as a terminal electron acceptor. As controls we also evaluated cells transformed with an empty vector and a native MtrA expression vector. After 24 hours (Figure 3a), all strains grew to a greater extent than the empty vector control with the exception of cells expressing IV-261. Additionally, cells expressing IV-301 grew to a significantly higher density than cells expressing native MtrA. A comparison of the z scores from the library analysis and OD_600_ from growth complementation revealed a positive linear relationship with an r^2^ = 0.93 (Figure S7). After 48 hours (Figure 3b), cells expressing IV-261 complemented growth. In contrast, cells transformed with an empty vector did not present growth even after 72 hours (Figure S8). All of the cultures with an OD_600_ increase also changed in color from brown to light yellow consistent with the reduction of Fe(III); this color change reversed upon exposure to oxygen. To investigate if random mutations beyond the peptide insertions led to the observed growth, vectors were sequenced following growth complementation. No mutations were observed beyond the peptide insertions. These findings show that the enrichment values observed in our library experiment correlate with growth complementation of individual variants. Additionally, they show that the variants having the lowest z scores support EET, albeit with severely decreased efficiencies compared with native MtrA.

**Figure 3.**
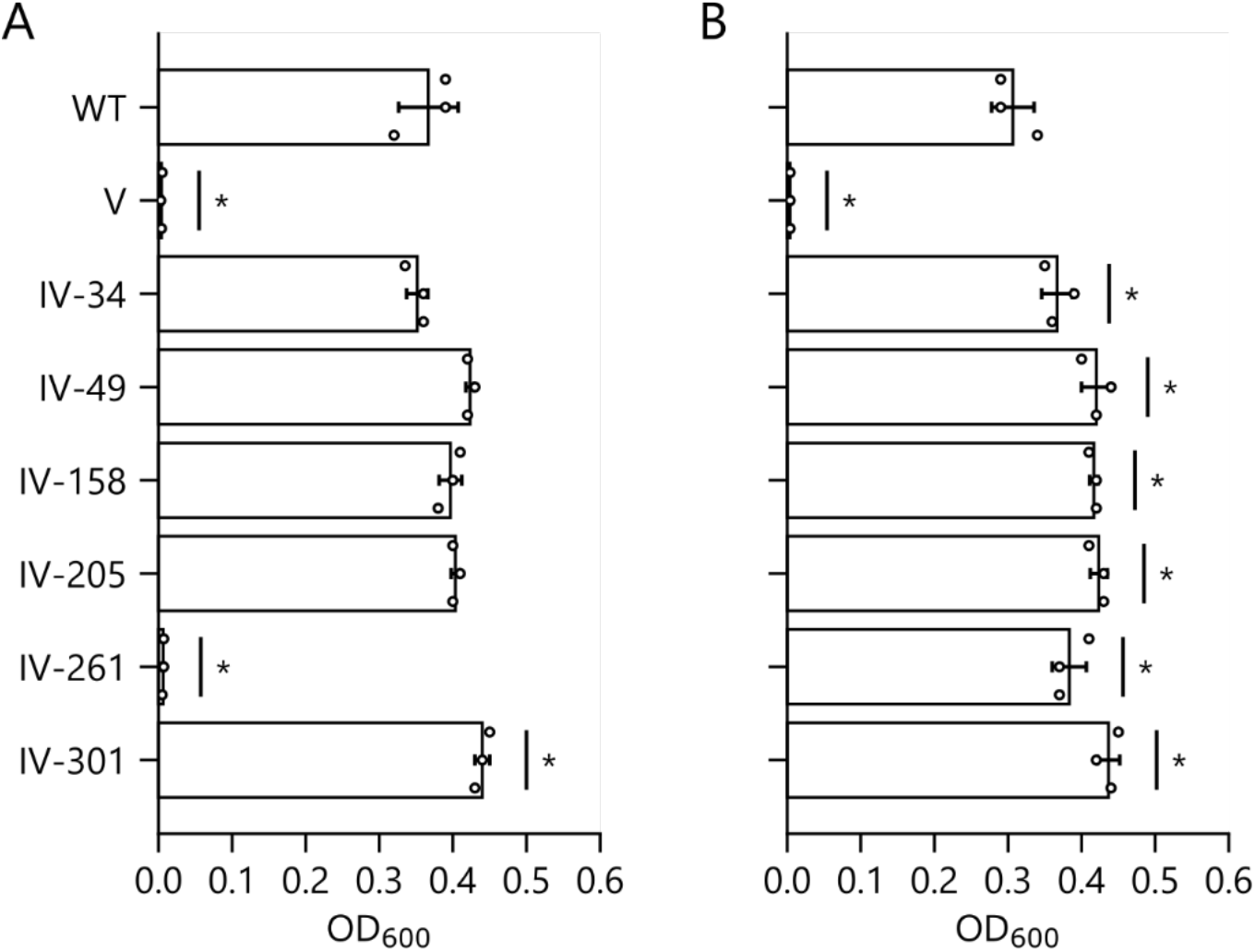
Complementation of *So*-*ΔmtrA* on ferric citrate by individual mutants. Cells transformed with an empty vector or plasmids that express wildtype and mutant MtrA were grown under anaerobic conditions with ferric citrate as the sole electron acceptor at 30°C. The optical density of cultures was measured at (**A**) 24 and (**B**) 48 hours. Error bars represent ±1σ from three experiments. After 24 hours, IV-301 grew to a significantly higher density than cells expressing native MtrA (P < 0.05 two-tailed t-test). All other mutants except IV-261 grew to a significantly higher density than the vector control (P < 0.01; two-tailed t-test).

**Table 1.** MtrA insertion variants targeted for characterization. For each variant, the table provides the name of the mutant, which is based on the native MtrA residue that follows the SGRPGSLS insertion, the identity of that amino acid that follows the insertion, the z score of the mutant from the library selection, and the plasmid name.

| **Name** | **Residue** | **Z score** | **Plasmid** |
| --- | --- | --- | --- |
| IV-34 | A34 | -2.52 | pIC014 |
| IV-49 | T49 | 5.05 | pIC017 |
| IV-103 | C103 | -4.14 | pIC032 |
| IV-158 | V158 | -0.003 | pIC015 |
| IV-165 | Q165 | -4.47 | pIC036 |
| IV-205 | A205 | 2.49 | pIC016 |
| IV-236 | C236 | -4.47 | pIC033 |
| IV-261 | P261 | -4.47 | pIC013 |
| IV-301 | N301 | 7.57 | pIC018 |

### Mutants with insertions in CXXCH motifs support ferric citrate reduction

Surprisingly, the variant depleted to the largest extent by the selection (IV-261) retained the ability to support respiration on Fe(III). For this reason, expression vectors for three additional variants (IV-103, IV-165, and IV-236) were created that present low z scores in the mutational profile (Table 1). Two of these variants, IV-103 and IV-236, were chosen because they insert a peptide within a CXXCH motif (Figure 4a), and are expected to prevent covalent ligation of hemes 2 and 7, respectively. All three variants were transformed into *So*-*ΔmtrA* and evaluated for growth complementation. After 24 hours, only cells expressing native MtrA showed significant growth (Figure 4b). After 72 hours, cells expressing all three variants presented optical densities comparable to cells expressing native MtrA and significantly higher than the empty vector control (Figure 4c). Cultures expressing these variants all displayed a color change characteristic of Fe(III) reduction. These findings show that MtrA variants having an octapeptide inserted within a CXXCH motif retain the ability to support EET. This tolerance is surprising because structural studies suggest that hemes 2 and 7 are required for rapid electron transfer across the length of MtrA.

**Figure 4.**
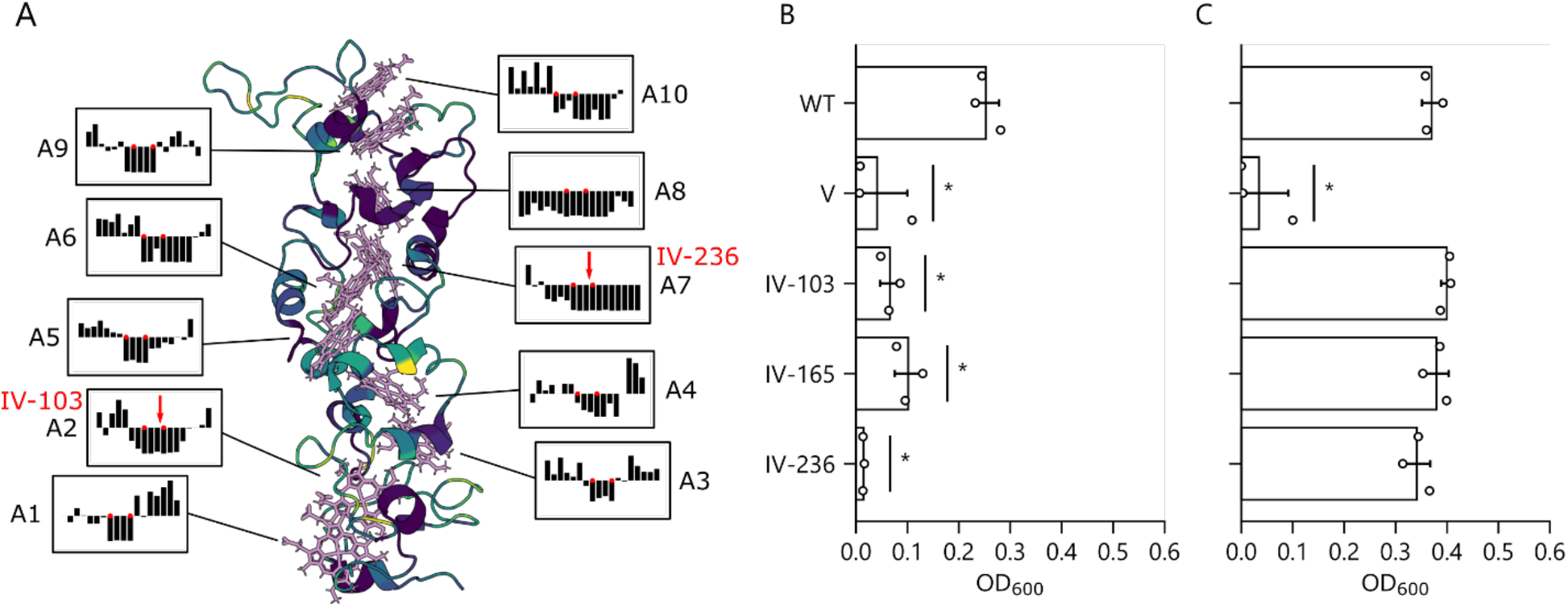
Effect of peptide insertion within CXXCH motifs on EET. (**A**) The MtrA structure is color coded using the z scores observed in the mutational profile, and insets depicting the distribution of z-scores around each of the 10 CXXCH motifs are shown. Cells transformed with an empty vector or plasmids that express wildtype and mutant MtrA were grown under anaerobic conditions with ferric citrate as the sole electron acceptor at 30°C. The optical density of cultures was measured at (**B**) 24 and (**C**) 72 hours. Error bars represent ±1σ from three experiments. After 72 hours, all tested IV’s grew to a significantly higher density than cells expressing vector control (P < 0.05 two-tailed t-test).

To determine if the octapeptide insertions decrease the amount of MtrA that accumulates as a decaheme holo-protein in cells, we measured the cytochrome c content of *So*-*ΔmtrA* cultures expressing all nine variants and native MtrA using enhanced chemiluminescence (ECL). Cells expressing MtrA presented bands consistent with the well-known cytochromes in *S. oneidensis*, including MtrC, MtrA, and CymA (Figure S9). In addition, two smaller cytochromes were observed that were interpreted as representing CcmE and NapB. Cells transformed with an empty vector contained all cytochromes except MtrA. In addition, the different insertion variants presented a range of MtrA levels. To account for loading variation, the intensities of MtrA bands were normalized to the band representing the smallest cytochrome, which was the most consistent cytochrome in our analysis (Figure 5a). When this normalization was performed, only a subset of the MtrA variants presented decreased levels, including two having insertions within a CXXCH motif (IV-103 and IV-236) and two with insertions outside of a heme-binding motif (IV-165 and IV-261). Surprisingly, all of these MtrA variants presented some holo-MtrA by ECL. The detection of holo-MtrA mutants having modified CXXCH motifs shows that the cytochrome c maturation system can covalently attach hemes to proteins with dramatic alteration to a single CXXCH motif. In case of the other variants that exhibited decreased holo-protein levels by ECL, this decrease could arise because the insertions occur in MtrA regions that are critical for interacting with the cytochrome c maturation system. Overall, these results show that peptide insertion within heme-binding motifs decrease the total holo-MtrA levels within cells, yet these mutants still mature into sufficient holo-MtrA to support cell growth that requires ferric citrate reduction.

**Figure 5.**
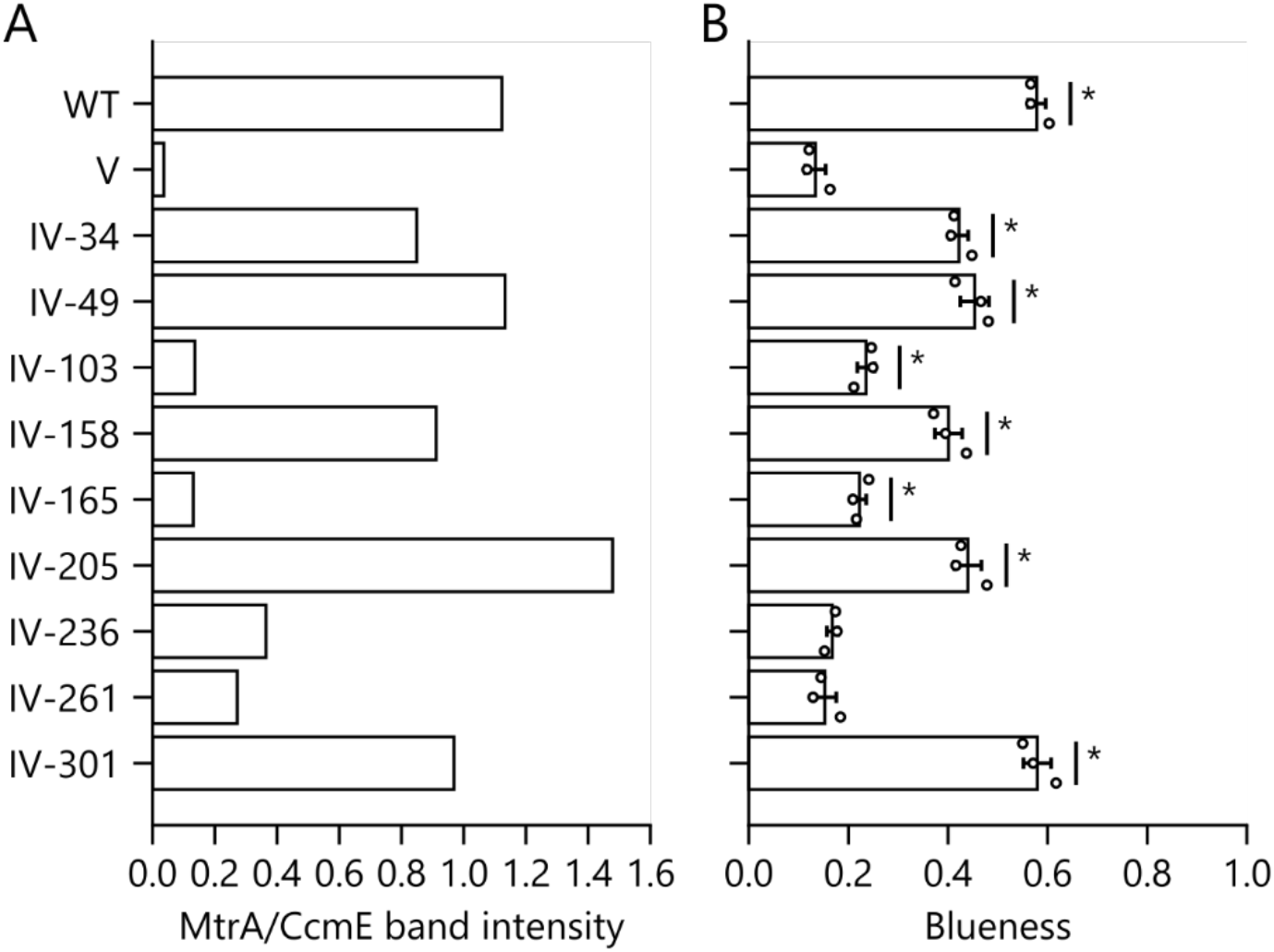
Effect of mutations on cytochrome maturation and EET to nanomaterials. *So*-*ΔmtrA* transformed with empty vector and plasmids expressing MtrA and nine different insertion variants were cultured overnight. Total protein from each sample was separated on an agarose gel and stained for cytochromes. (**A**) Ratio of MtrA/CctA band intensities for ECL samples. (**B**) *So*-*ΔmtrA* taken from the same culture as the ECL samples were incubated in M9 minimal medium containing 10 mg/L MoO_3_ nanoparticles under anaerobic conditions for 15 minutes in a 96-well plate. The blueness of each well was quantified. Samples presenting significantly higher signal compared with the empty vector are noted with an asterisk (P < 0.05; two-tailed t-test). Error bars represent ±1 standard deviation calculated from three experiments.

### Mutants differentially reduce soluble and insoluble electron acceptors

MtrA transfers electrons to the surface-displayed cytochrome MtrC, which in turn modulates binding^14^ and electron transfer to both soluble Fe(III) and insoluble extracellular materials.^42^ These observations led us to hypothesize that all of our variants with decreased EET to Fe(III) would also present decreased EET to materials. To determine the effects of MtrA mutations on electron transfer to extracellular materials, we utilized a modified version of a nanoparticle reduction assay.^43^ In this assay, we evaluated EET to MoO_3_ nanoparticles (Figure S10), rather than commonly used WO_3_, since we found that MoO_3_ particles are more sensitive for reporting on EET.^44^ With this assay, the nanoparticles undergo an electrochromic shift from white to blue as electroactive microbes reduce them via EET. When we used this assay to characterize *So*-*ΔmtrA* cells expressing different insertion variants (Figures 5b; S11), we found that only a subset of the variants support MoO_3_ reduction following a 15 minute incubation. Cells expressing insertion variants with positive z scores showed rapid and strong reduction of the nanoparticles, with IV-301 (z = 7.57) matching the reduction achieved with native MtrA. More than half of the variants with negative z scores (IV-103, IV-165, and IV-236) supported nanoparticle reduction that significantly exceeded the vector control, although they showed lower reduction capabilities than variants with positive z scores. Two variants (IV-34 and IV-261) were unable to reduce the nanoparticles beyond the negative control, even after a 96 hour incubation. This finding indicates that cells expressing some insertion variants failed to reduce MoO_3_ even though they were competent at reducing extracellular ferric citrate. Among the insertion variants, those that presented the greatest reduction in the nanoparticle assay had the largest abundance of holo-MtrA as judged by ratios of MtrA/CcmE in the ECL analysis. In contrast, those variants with low or no signals in the nanoparticle assay exhibited faint bands at the same position. Interestingly, IV-236 showed a stronger MtrA/CcmE ratio than IV-103 and IV-165, both of which outperformed IV-236 in the nanoparticle assay.

To probe the effect of MtrA peptide insertion on EET relevant for microbial electrochemical technologies, we evaluated the ability of *So*-*ΔmtrA* strains expressing MtrA variants to reduce an electrode using chronoamperometry. We evaluated the current generated by cells expressing native MtrA and variants exhibiting a range of EET with ferric citrate and MoO_3_ nanoparticles, including the most active variant (IV-301), least active variant (IV-261), and two with intermediate z scores (IV-205 and IV-103) (Figure 6a). We hypothesized that the current observed in a bioelectrochemical system would correlate more strongly with the signal obtained with MoO_3_ nanoparticles. As controls, we also evaluated the current generated by *So*-*ΔmtrA* transformed with an empty vector and wildtype *S. oneidensis* MR-1. With cells expressing IV-261 and IV-103 (z scores = -4.14 and -4.47), current production was indistinguishable from the empty vector control. In contrast, cells expressing native MtrA and IV-301 (z score = 7.57) presented similar current densities. In contrast, IV-205 (z score = 2.40) presented an intermediate current density, transferring a total charge significantly greater than the empty vector control yet significantly lower than the strain expressing native MtrA (Figure 6b). For the cells expressing insertion variants, the total charge transferred over the course of the experiment was linearly correlated with the z score observed in the library selection (r^2^ = 0.96) on ferric citrate (Figure 6c). Like *S. oneidensis* MR-1, *So*-*ΔmtrA* expressing IV-301, IV-205, and MtrA showed strong initial current peaks that diminished within 5 hours. These findings reveal a striking correlation between the sequence enrichments arising from the selection with a soluble electron acceptor and the current generated at an electrode. Materials like electrodes can be challenging to directly use in library selection as changes in electron flux can be confounded by changes in biofilm formation or surface interactions.^45^

**Figure 6.**
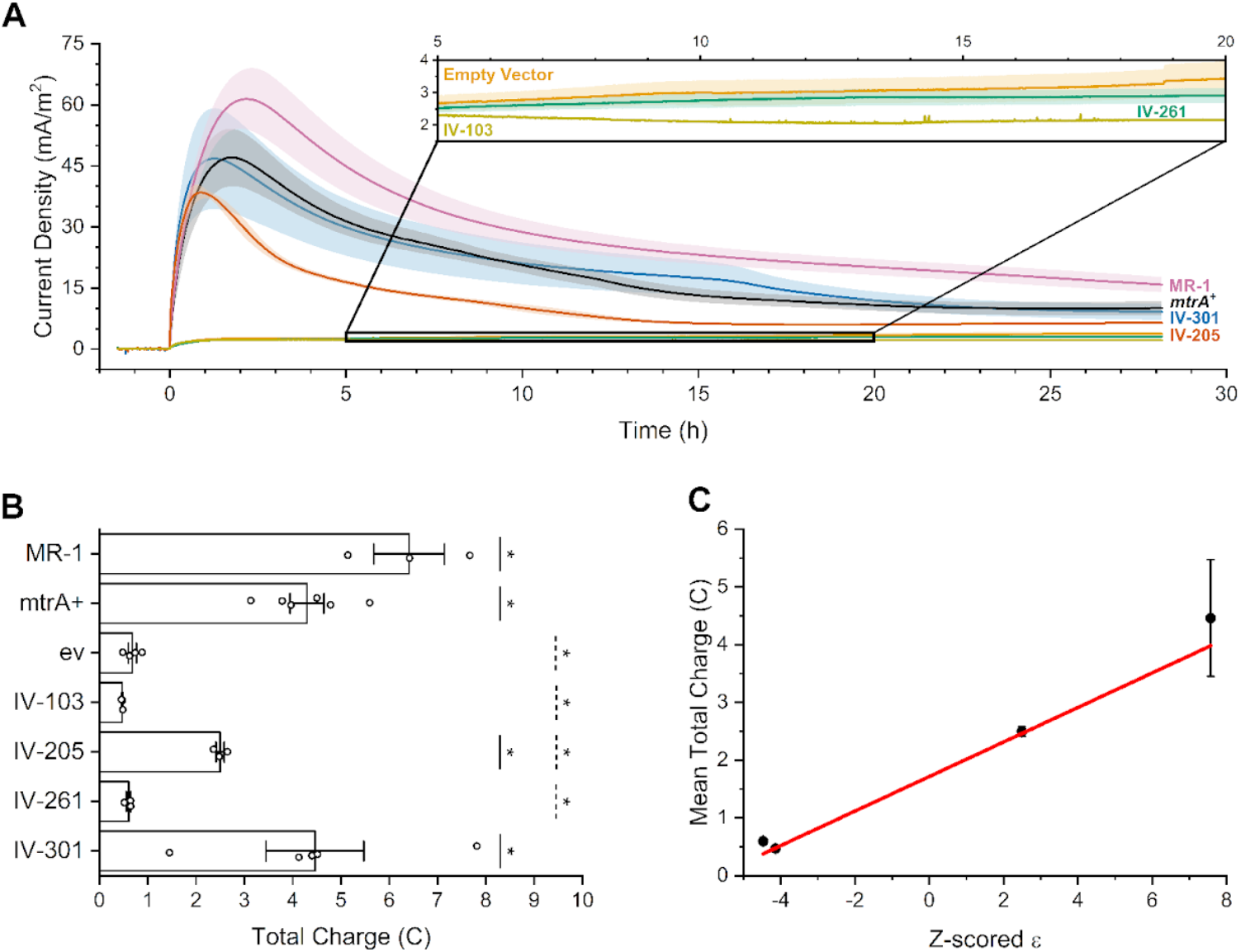
Effect of mutations on current in a bioelectrochemical cell. (**A**) Chronoamperometry of *So*-*ΔmtrA* expressing native MtrA (black), IV-261 (green), IV-301 (blue), IV-103 (yellow), IV-205 (red), and empty vector (orange) are compared with *Shewanella oneidensis* MR-1 (purple). *So*-*ΔmtrA* expressing IV-103 and IV-261 produce negligible current, while *So*-*ΔmtrA* expressing IV-205 and IV-301 approach native MtrA values. Experiments were performed using a carbon felt working electrode at +0.2 V vs. Ag/AgCl (3 M KCl) with 30 mM D,L-lactate as the electron donor. Data are plotted as the mean of biological replicates for MR-1 (n = 3) and *So*-*ΔmtrA* expressing MtrA (n = 6), IV-103 (n = 3), IV-205 (n = 3), IV-261 (n = 3), IV-301 (n = 5), and *So*-*ΔmtrA* transformed with an empty vector (n = 4). Error bars represent ±1 standard error of the mean. Time is listed relative to the injection of cells into the bioelectrochemical reactor. (**B**) Total charge transferred to the electrode by each strain, computed by integrating the baseline subtracted current over the period from t = 0 hours to experiment end. Strains that transferred significantly more charge compared with the empty vector are noted with a solid line and an asterisk (P < 0.05; two-tailed t-test). Strains that transferred significantly less charge compared with *So*-*ΔmtrA* expressing native MtrA are noted with a dashed line and an asterisk (P < 0.05; two-tailed t-test). Error bars represent ±1 standard error of the mean. (**C**) Total charge transferred by *So*-*ΔmtrA* expressing IV-261, IV-301, IV-103, and IV-205 compared against the z scores observed in the mutational profile for these insertion variants, with a linear regression fitted to the data (r^2^ = 0.96). Error bars represent ±1 standard error of the mean.

### Insertion tolerance correlates with heme proximity

To evaluate the structural underpinnings for MtrA mutant activity, we mapped MtrA mutant z scores onto the *Shewanella baltica* MtrA structure.^20^ Regions that led to the largest decreases in EET following peptide insertion included: (i) the loops at the interface between adjacent perpendicular hemes, (ii) a loop which interacts with the MtrB pore close to the extracellular surface of the lipid bilayer, (iii) residues that are localized to the periplasm, and (iv) all ten alpha-helices coordinating the ten heme cofactors (Figure 2). Insertion tolerance is low within all ten CXXCH heme-ligating motifs, but it is not negligible and insertions are tolerated within the regions flanking these motifs. Unexpectedly, the interface between MtrA and MtrC shows a high level of peptide insertion tolerance. Thus, our systematic mutagenesis approach identifies diverse regions and motifs that tolerate peptide insertion without disrupting EET.

To enable efficient ET across the outer membrane, MtrA must arrange ten iron-containing heme prosthetic groups at optimal distances and orientations for electron transfer within a protein environment. To investigate if peptide insertion tolerance depends upon proximity to heme prosthetic groups, we evaluated how insertion tolerance varies with proximity to the central axis of MtrA containing the heme prosthetic groups and position along the long axis of MtrA that spans the outer membrane. Consistent with the idea that MtrA functions as a protein wire, a correlation was observed between sequence enrichment values (ε_i_) and radial insertion distance from the center axis of iron atoms (r_s_ = 0.48) (Figure 7a). In addition, a positive correlation was observed between insertion tolerance and the sequence distance to the nearest heme-ligated cysteine (r_s_ = 0.35) (Figure 7b). In contrast, no correlation was observed between insertion tolerance and position along the long axis of MtrA (Figure 7c), and a weak correlation (r_s_ = -0.39) was observed between insertion tolerance and the density of intramolecular residue-residue contacts (Figure 7d) when both short- and long-range contacts (≤14 Å) were considered. No correlation was observed when comparing insertion tolerance to short range (≤8 Å) contacts (Figure S12a) and the density of interprotein residue-residue contacts between MtrA and MtrB (Figures S12b). A majority of the MtrA and MtrC residue-residue contacts presented enrichment, with a weak positive correlation observed between enrichment and number of interprotein contacts between MtrA and MtrC (Figures S12c). No correlation was observed with B factors from the MtrA crystal structure (Figure S12d) and stiffness calculated from a coarse-grain elastic network model (Figures S12e). Together, these analyses show that the strongest correlations are observed between insertion tolerance and proximity to the axis of iron atoms that span the outer membrane and function as a wire, proximity to CXXCH motifs through primary structure, and the density of intramolecular residue-residue contacts.

**Figure 7.**
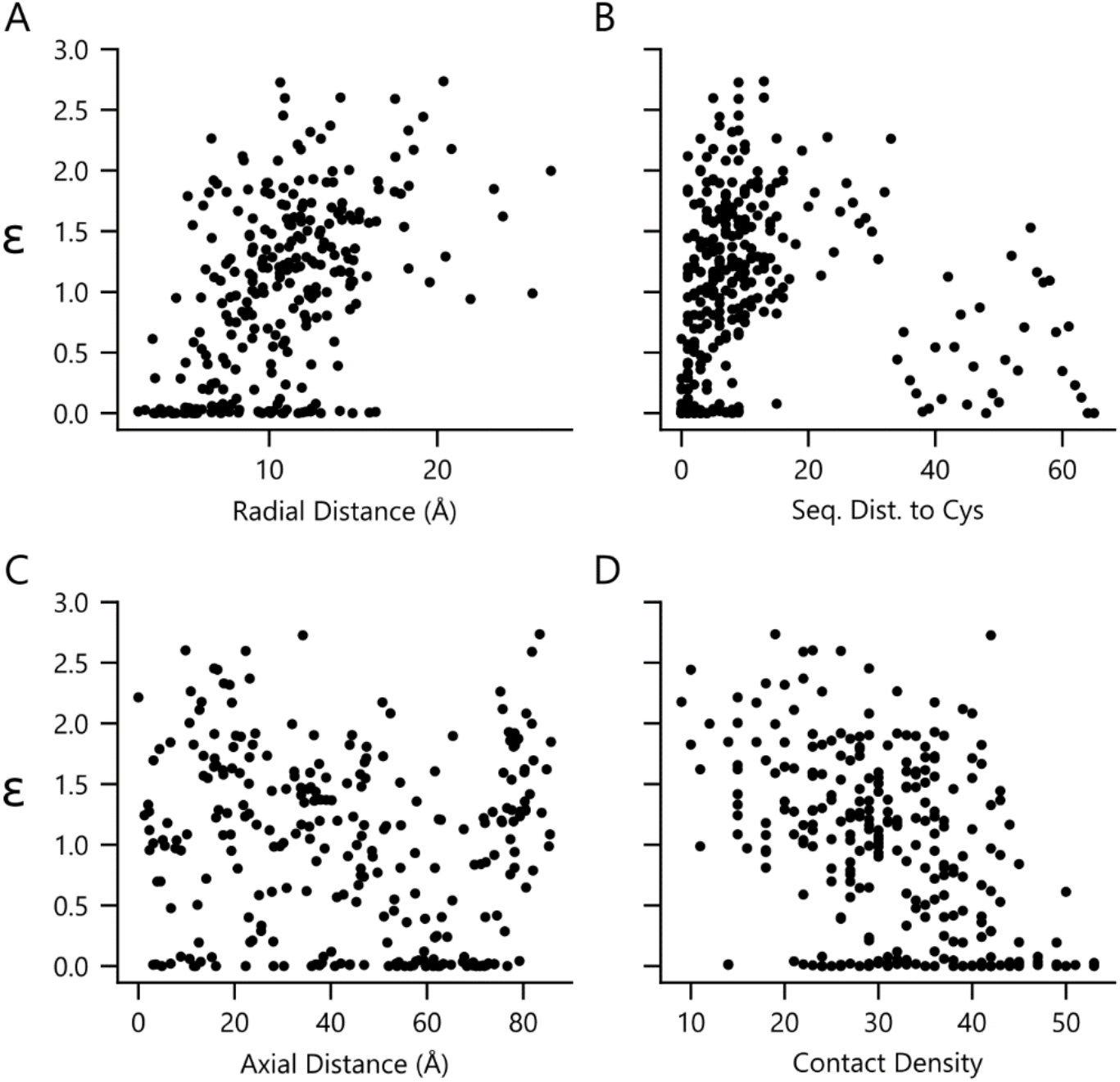
Comparison of insertional tolerance and structural features in MtrA. Spearman correlations between these z scores and biophysical properties were evaluated, including: (**A**) radial distance from a central axis defined by heme iron atoms (r_s_ = 0.48; p < 10^-15^), (**B**) distance in primary structure from heme-binding cysteines (r_s_ = 0.35; p < 10^-10^), (**C**) axial distance (r_s_ = -0.10; p = 0.1), and (**D**) intramolecular MtrA contacts ≤14 Å (r_s_ = -0.39; p < 10^-10^).

## DISCUSSION

Our results greatly extend our fundamental understanding of sequence-structure-EET relationships in multiheme cytochromes. Prior genetic studies revealed the functional roles that different members of the MtrCAB complex play in supporting EET to extracellular metal ions and materials.^15–17, 22, 46, 47^ However, prior to this study, only two studies examined the effect of mutations on MtrA. The first study demonstrated that the histidines within the MtrA CXXCH motifs are critical for EET,^48^ while the second study identified a pair of mutations in MtrA and MtrB that compensate for the loss of MtrC under anaerobic conditions.^49^ Additionally, only two studies evaluated the effects of mutations on MtrC-type multiheme cytochromes, which transfer electrons to insoluble electron acceptors. These efforts revealed that the cysteines in the MtrC CX_8_C motif form a disulfide that is critical for aerobic growth^50^ and identified residues that modulate nanoparticle binding through electrostatic interactions.^14^ By using a combinatorial mutagenesis approach to comprehensively evaluate the functional consequences of generating mutations at every native location in a multiheme cytochrome, we identify diverse regions across the primary structure that present a high functional tolerance to peptide insertion, such as extracellular residues proximal to the MtrA-MtrC binding interface. Additionally, we establish the motifs that are most sensitive to peptide insertion, such as the CXXCH motifs. Surprisingly, Fe(III) reduction was still observed following CXXCH disruption with peptide insertions, a mutation that is more structurally disruptive than previously studied point mutations.^48^ This finding suggests that individual heme binding sites can be structurally disrupted without completely abolishing MtrA maturation, insertion into MtrB and the outer membrane, and MtrA-mediated EET.

Our results provide insight into residue-residue contacts made by MtrA and MtrB that are critical to supporting EET. Although the mechanism of insertion of MtrA into MtrB is not well understood, it is clear that MtrA protects MtrB from degradation, enabling expression of MtrB.^51^ Indeed, the MtrB channel is narrower at the extracellular end compared with the periplasmic end, leading to prior speculation that MtrB arrests MtrA in state of stalled excretion.^20^ The interface between MtrB and MtrA is not thought to be tight, although a solvent channel has been proposed to run between MtrB and one side of MtrA, with residues from both proteins capping this channel.^20^ Our measurements revealed variation in the insertion tolerance proximal to MtrA residues that make contacts with MtrB. Among the diverse positions that make contact with MtrB, several showed a high tolerance to insertion, especially those closer to the periplasmic end of the channel. In contrast, some regions closer to the extracellular end of the channel presented marked intolerance to peptide insertion. In particular, the loop defined by MtrA residues 240-247 had a large number of MtrB contacts that exhibited a high functional sensitivity to peptide insertion. These findings implicate this interacting motif as critical to MtrAB secretion and/or stability.

Surprisingly, the interface between MtrA and MtrC did not appear sensitive to peptide insertion in our library selection. In fact, the peptide insertion site enriched to the greatest extent is located in an extracellular-facing loop at the MtrA-MtrC interface. The underlying mechanism by which MtrA insertion variants retain function upon peptide insertion at the MtrA-MtrC interface is not known. Prior structural studies have provided evidence that MtrA and MtrC use hydrogen bonds to position themselves closely together, with heme A10 of MtrA being only 8 Å from heme C5 of MtrC.^20^ Additionally, genetic studies have shown that native MtrA cannot reduce ferric citrate above detectable limits in the absence of MtrC.^22, 47^ One possible explanation for the high EET observed with peptide insertion near the MtrA-MtrC binding interface is that the mutant MtrA-MtrB complexes exhibit enhanced ferric citrate reduction activity. A previous study showed that mutations in MtrA and MtrB increase ferric citrate reduction in a *S. oneidensis* strain lacking MtrC and its homologs.^49^ Whether MtrA peptide insertions allow the MtrA-MtrB complex to reduce ferric citrate will require future studies of EET in MtrC knockout strains.

The observation that some insertion variants vary in their ability to reduce soluble (ferric citrate) and insoluble (nanomaterials and electrode) electron acceptors suggests that some insertion variants may not require MtrC to respire. In a prior study, a mutated MtrA-MtrB complex could reduce soluble extracellular acceptors in the absence of MtrC, such as 9,10-Anthraquinone-2,7-disulphonic acid (AQDS) and ferric citrate.^49^ However, these mutants were diminished in their ability to reduce insoluble extracellular acceptors like birnessite and pahokee peat humic acids.^49^ This observation highlights the role that MtrC plays in facilitating EET to larger terminal electron acceptors. Taken together with our results, this finding suggests that MtrA insertion variants that retain the ability to reduce ferric citrate but fail to reduce insoluble extracellular electron acceptors may be operating in an MtrC-independent manner. This trend is expected to occur when peptide insertion disrupts the interaction between heme A10 in MtrA and from heme C5 in MtrC, which could arise from steric clashes caused by peptide insertion or conformational changes in MtrA induced by peptide insertions. ET between these hemes is thought to be the slowest within MtrCAB complex and a subtle structural change in one of these prosthetic groups could disrupt ET from MtrA to MtrC, cutting off MtrC from the rest of the pathway.^4^

The N terminus of MtrA, which is unresolved in the MtrA crystal structure,^20^ exhibited a low tolerance to peptide insertion. This region does not ligate a heme so this trend is not thought to arise from disrupting MtrA ET. This trend could arise through three different mechanisms. First, the initial thirty-four residues in MtrA comprise the signal peptide sequence responsible for translocation across the inner membrane.^42^ Disruption of this motif could impede interaction with the Sec translocase and decrease the amount of MtrA that enters the periplasm.^52^ Second, peptide insertion near the N terminus could decrease MtrA expression due to proximity to the RBS that regulates translation initiation. A thermodynamic model has revealed that mutations proximal to an RBS can generate local RNA structures that compete with ribosome binding.^53^ Third, peptide insertion could disrupt the region of MtrA that mediates binding to periplasmic electron carriers. Genetic and structural studies have implicated several periplasmic and inner membrane-bound oxidoreductases in reducing MtrA, including CymA, CctA, and FccA.^22, 54, 55^ Given that CctA is knocked out in the strain we utilized for our selection assays (JG665), the sensitivity of MtrA to insertions at the N terminus suggests this region mediates contact with CymA or FccA.^22^ In the future, it will be interesting to select MtrA mutant libraries in strains that contain different periplasmic electron carriers and to compare the results to determine if residues in the N-terminal region of MtrA present a similar functional sensitivity to peptide insertion.

Our results suggest that the underlying mechanism by which peptide insertion disrupts EET may vary from mutant to mutant. For example, ECL measurements of holo-MtrA suggest that the decreased EET to MoO_3_ nanoparticles may arise because the total amount of holoprotein that accumulates in the membrane is diminished. This appears to occur with IV-165 and IV-261, which present both decreased EET to MoO_3_ nanoparticles and lowered MtrA/CcmE ratios. However, other insertion variants, like of IV-236 showed a stronger MtrA/CcmE ratio than IV-165, yet it presented weaker EET in the nanoparticle assay. This finding suggests that peptide insertion in IV-236 has little effect on holo-MtrA levels but instead disrupts EET by disrupting the distances and orientations of a heme prosthetic group or by altering the binding affinity for partner oxidoreductases like MtrC and periplasmic carrier partners.

The library approach herein represents an excellent starting point for further probing sequence-structure-EET relationships in multiheme cytochromes that mediate EET. Similar libraries can be generated within other members of the MtrCAB complex to better understand critical residues for their molecular interactions. Additionally, larger insertions can be introduced into these proteins to better understand how they can be engineered for bioelectronics applications.^56^ In future studies, it will also be interesting to investigate whether ligand binding domains that exhibit allosteric conformation changes can be inserted into MtrA and used to regulate EET through ligand-binding events, similar to that previously achieved with cytosolic protein electron carriers.^57, 58^ Allosteric protein switches that control EET on the extracellular face of cells would have distinct advantages over cytosolic protein switches because they would be regulated by analyte concentrations in the extracellular environment, which can be distinct from the concentrations that enter the cell.

## METHODS

### Strains

*E. coli* XL1-Blue was from Agilent, and *dam-/dcm-E. coli* was obtained from New England Biolabs. *S. oneidensis* JG665 (*So*-*ΔmtrA*) was a kind gift from Dr. Jeff Gralnick,^22^ while *S. oneidensis* MR-1 was from the American Type Culture Collection.

### Plasmid construction

Plasmids are listed in Table S1. Endogenous genes were synthesized as G-blocks by Integrated DNA technologies, and plasmids were constructed by using Q5 High-Fidelity DNA polymerase (New England Biolabs) to produce amplicons and ligating amplicons to PCR amplified vectors using Golden Gate DNA assembly.^59^ The broad range RSF1010 backbone was used for all plasmids and was given generously by Shyam Bhakta.^60^ All plasmid sequences were verified with Sanger sequencing.

### Library generation

The MtrA protein (UniProt Q8EG35) was used for library creation. Oligonucleotides encoding MtrA gene fragments with insertions encoding SGRPGSLS were designed using the SPINE algorithm.^28^ Gene tiles synthesized as an oligo pool by Twist Bioscience were PCR amplified using Q5 High-Fidelity DNA polymerase and cloned into the vector RSF1010 using Golden Gate cloning. Plasmid ensembles were transformed into *E. coli* XL1-Blue and plated onto lysogeny broth (LB) agar plates containing 50 µg/mL kanamycin. The transformations for each tile yielded >2500 colony forming units. Assuming sampling with replacement, this colony count indicates that >99% of our variants were sampled at this step.^37^ Colonies were harvested from plates and pooled, and plasmid DNA was purified using a Qiagen Miniprep kit. The distribution of insertion sites was evaluated using Amplicon-EZ sequencing (Genewiz). The library was then transformed into *dam-/dcm-E. coli* (New England Biolabs), and sufficient colony counts were obtained to sample >99% of our variants. The demethylated insertion library was transformed into *S. oneidensis So*-*ΔmtrA* using electroporation and plated on LB agar plates containing 50 µg/mL kanamycin. The naïve library harvested from these plates was stored as a glycerol stock.

### Growth complementation

*So*-*ΔmtrA* transformed with plasmids expressing *mtrA* or empty vector were grown alongside *S. oneidensis* MR-1 transformed with a vector bearing *mtrA* overnight at 30°C and 250 rpm in Shewanella Basal Medium (SBM) containing 8.6 mM NH_4_Cl, 1.3 mM K_2_HPO_4_, 1.65 mM KH_2_PO_4_, 475 µM MgSO_4_·7H_2_O, 1.7 mM (NH_4_)_2_SO_4_, 7.5 mg/L nitrilotriacetic acid, 0.5 mg/L MnCl_2_·4H_2_O, 1.5 mg/L FeSO_4_·7H_2_O, 0.85 mg/L CoCl_2_·6H_2_O, 0.5 mg/L ZnCl_2_, 0.2 mg/L CuSO_4_·5H_2_O, 0.025 mg/L AlK(SO4)_2_·12H_2_O, 0.025 mg/L H_3_BO_3_, 0.45 mg/L Na_2_MoO_4_, 0.6 mg/L anhydrous NiCl_2_, 0.1 mg/L NaWO_4_·2H_2_O, 0.5 mg/L Na_2_SeO_4_, 0.01 mg/L biotin, 0.01 mg/L folic acid, 0.1 mg/L pyridoxine HCl, 0.025 mg/L thiamine, 0.025 mg/L nicotinic acid, 0.025 mg/L pantothenic acid, 0.5 µg/L cyanocobalamin, 0.025 mg/L p-aminobenzoic acid, 0.025 mg/L alpha-lipoic acid, 100 mM HEPES (pH 7.2), 0.05% (w/v) casamino acids, 20 mM lactate, and 50 µg/mL kanamycin.^61^ Overnight cultures were diluted 1:100 in fresh SBM supplemented either with 30 mM ferric citrate or 30 mM fumarate and transferred to a 96 well plate. The plate was incubated in a N_2_ atmospheric chamber at 30°C and shaken at 500 rpm. Optical density measurements at 600 nm (OD_600_) were taken after 48 hours using a Tecan Spark plate reader.

### Library selections

The glycerol stock of the naïve library was used to inoculate SBM medium which were grown for 18 hours aerobically at 30°C while shaking at 250 rpm. To evaluate the sequence diversity in the naive library, three separate overnight cultures were harvested and subjected to deep sequencing (Genewiz). An aliquot of each culture was also diluted into fresh SBM medium to an OD_600_ = 0.1 and used to inoculate sealed bottles containing either SBM supplemented with 30 mM fumarate (n = 3) or SBM supplemented with 30 mM ferric citrate (n = 3). This growth medium was sparged for 30 minutes with N_2_ prior to inoculation, and it was filtered to remove all undissolved ferric citrate. After incubation for 24 hours, cells were harvested and samples were sent for deep sequencing.

### Sequence analysis

Table S2 provides the relative abundances of each variant observed in the three different naive, unselected, and selected libraries. Variants were numbered using the native residue that immediately follows the peptide insertion, *e.g.*, IV-261 represents the variant having a peptide inserted before Pro261. Insertion sites identified from deep sequencing were processed using the DIP-seq algorithm.^62^ Two sequence enrichments were calculated for each variant: enrichment after the selective growth in ferric citrate relative to the naïve condition (ε_s_) and enrichment after non-selective growth in fumarate relative to the naïve condition (ε_n_). Enrichments were calculated by dividing the frequency (f_i_) of each variant observed in ferric citrate (f_s_) or fumarate (f_n_) conditions by the frequency of each variant in the naïve library. The frequencies of the naïve and non-selective insertion libraries were then fit to a linear regression using NumPy.^63^ A z-score for ε_s_ values was generated by subtracting the mean of the ε_n_ distribution (µ_n_) and normalizing by the standard deviation of the ε_n_ distribution (σ_n_). Z-scored ε_s_ were overlaid onto the structure of MtrA from *Shewanella baltica* (PDB: 6r2q)^20^ and visualized using PyMol (Schrodinger, LLC. 2010. The PyMOL Molecular Graphics System, Version 2.5).

### MoO_3_ nanoparticle assay

Cultures grown for 16 hours in LB containing 50 µg/mL kanamycin were pelleted using centrifugation (4000 x g) for 10 minutes at room temperature, washed with an M9 medium containing sodium phosphate, dibasic (6.8 g/L), sodium phosphate, monobasic (3 g/L), sodium chloride (0.5 g/L), 2% glucose, ammonium chloride (1 g/L), and 20 mM lactate, and resuspended to an OD_600_ of 1. Resuspended cultures were mixed at a 1:1 ratio with 10 mg/mL molybdenum trioxide (MoO_3_) nanoparticles (Sigma-Aldrich) and incubated for 15 minutes in a 96-well plate at room temperature in a 5% H_2_, 10% CO_2_, and 85% N_2_ atmosphere. Cultures were scanned using a CanoScan LiDE 220 and the blueness of wells was determined using ImageJ with the ReadPlate3 plugin.^64^ The comparison experiment between MoO_3_ and WO_3_ was conducted in the same manner except that precultures were grown to an OD_600_ = 5 and incubated for 10 minutes; for these measurements we only compared *S. oneidensis* MR-1 EET.

### Enhanced chemiluminescence staining

Cultures grown for 16 hours in LB containing kanamycin (50 µg/mL) were pelleted (4000 x g) for 10 minutes at room temperature and washed with M9 medium as in the MoO_3_ assay. Cultures mixed in a 1:1 ratio with a gel loading solution containing mercaptoethanol (0.71 M) and 1X LDS Loading Dye Mix (NuPage) were incubated for 15 minutes. Samples were then loaded onto a 12% Bis-Tris gel (Invitrogen) and run for 20 min at 200 mV with a Precision Plus Protein Dual Color Standards ladder. Protein samples were transferred to blotting paper using a Trans-Blot Turbo Transfer Pack (Bio-Rad) and a Trans-Blot Turbo (Bio-Rad) using the “1-mini gel” setting. Cytochromes were visualized using Clarity Western ECL Substrate (Bio-Rad) then imaged for both visible and chemiluminescent signals using a FluorChem E imager (ProteinSimple). Chemiluminescent images were overlaid on visible images to compare protein standards and chemiluminescent cytochromes.

### Chronoamperometry

Each biological replicate was derived from a different colony picked from an LB-agar plate (with 50 µg/mL kanamycin) streaked with cells from a glycerol stock. Picked colonies were used to inoculate 5 mL cultures of LB with 50 µg/mL kanamycin that were grown for 16-20 hours at 30 °C, shaking at 250 rpm. These cultures were then used to inoculate 50 mL LB cultures (with 50 µg/mL kanamycin) in 250 mL flasks to a standardized OD_600_ of 0.01. The resulting 50 mL cultures were grown for 20 hours at 30 °C, shaking at 250 rpm. Cells were pelleted via centrifugation and the LB was removed. The cell pellets were washed twice with 25 mL of M9 minimal salts (BD). The anodic chambers of dual-chamber bioelectrochemical reactors were inoculated to a standardized OD_600_ of 0.5 using 3 mL of cells resuspended in M9 minimal salts. The anodic chamber contained a carbon felt working electrode (geometric surface area = 0.0022 m^2^, Alfa Aesar) suspended from a 1 mm diameter titanium wire, a Ag/AgCl (3 M KCl, CHI111, CH Instruments) reference electrode, and 117 mL of M9 minimal salts with 30 mM D,L-lactate (Sigma). This chamber was separated by a cation exchange membrane (CMI-7000, Membranes International) from a cathodic chamber containing 115 mL of M9 minimal salts and a 1 mm diameter titanium wire counter electrode. All electrodes were connected to a BioLogic VMP-300 potentiostat. The anodic chamber was continuously stirred with a stir bar at 300 rpm, temperature-controlled at 30 °C, and sparged with N_2_ gas to maintain anaerobic conditions. The working electrode was poised at +0.2 V versus Ag/AgCl and the average current was recorded every 36 seconds.

### Structural analysis and modeling

The crystal structure of the *Shewanella baltica* MtrCAB complex (PDB ID, 6R2Qu was used for analysis.^20^ Unresolved loops near the N-terminus of MtrA were reconstructed using Rosetta’s loopmodeler application on default settings using the beta_nov16-genpot forcefield,^65^ with MtrB, MtrC, and the heme prosthetic groups included. The lowest scoring of 500 structures from Rosetta was used for all further analysis unless otherwise specified. The axial position of each residue was calculated as distance from the Cα atom of the residue to the x-z plane normal to the long dimension of MtrA and centered at the Cα atom of lysine 62, the most intracellular residue resolved in the crystal structure. The radial position of each residue was calculated as distance from the Cα atom of the residue to the center axis defined by the average x and z coordinates of the iron atoms of all hemes in MtrA. To determine per-residue contact density, short-range contacts were identified between residues with a cutoff of 8 Å between Cα atoms, and both short- and long-range contacts were identified with a cutoff of 14 Å, as commonly used.^66–68^ Contacts were counted separately for residues within MtrA as well as between MtrA and MtrB and between MtrA and MtrC. Cα B factors were obtained from the crystal structure for each residue in MtrA using the Biopython Bio.PDB package^69, 70^ and averaged for the residue preceding and the residue following each insertion site. Per-residue stiffness estimates were obtained from a Cα anisotropic network model of the entire MtrCAB complex (without hemes) using the ProDy suite,^71, 72^ based on previously described protocols.^68, 73^

### Statistics

All error bars represent ±1 standard deviation calculated from three or more biological replicates. Independent, two-tailed t-tests were used to compare differences between all relevant samples with α = 0.05. All correlations shown are Spearman correlations calculated with Pandas.^74^

## ACCESS CODES

MtrA: Q8EG35

MtrB: Q8CVD4

MtrC: Q8EG34

## ACKNOWLEDGEMENTS

Funding was provided by Office of Naval Research grant N00014-20-1-2274 (to CMAF and JJS); Office of Science, Office of Basic Energy Sciences of the U.S. Department of Energy grant DE-SC0014462 (to JJS); Cancer Prevention and Research Institute of Texas RR190063 (to CMAF); and National Science Foundation (NSF) grant 1843556 (to JJS). MDC is supported by NSF National Research Traineeship in Bioelectronics grant 1828869, and JTA is supported by a NSF postdoctoral fellowship. Modeling was supported by Rice University’s Center for Research Computing and the Big-Data Private-Cloud Research Cyberinfrastructure NSF MRI-award (1338099). *S. oneidensis* JG665 (*So*-*ΔmtrA*) was a kind gift from Dr. Jeff Gralnick.

