## Supplemental Data for "Determinants of multiheme cytochrome extracellular electron transfer uncovered by systematic peptide insertion"

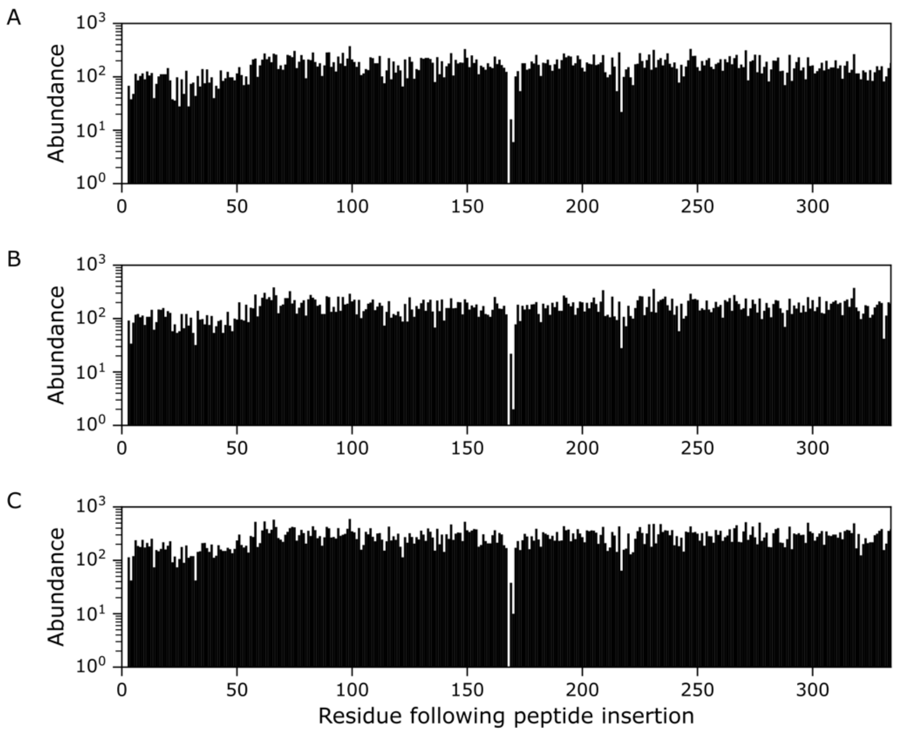


**Figure S1. Naive library sequence abundances in each experiment.** Sequencing data was filtered for reads that contain peptide insertions within the MtrA sequence. The abundance of reads for each insertion variant is listed for: (**A**) replicate 1 (n = 53148 filtered reads), (**B**) replicate 2 (n = 53824), and (**C**) replicate 3 (n = 90494).

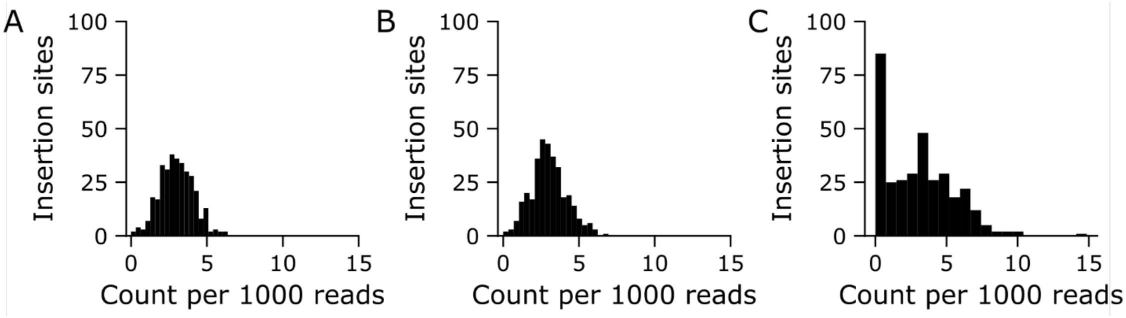


**Figure S2. Comparison of the frequency distributions for each library.** The distribution of insertion frequencies in (**A**) the naïve library (CV = 0.37) and (**B**) non-selected library (CV = 0.39) present similar CVs (0.37 and 0.39, respectively). (**C**) In contrast, the selected library presents a higher CV = 0.82.

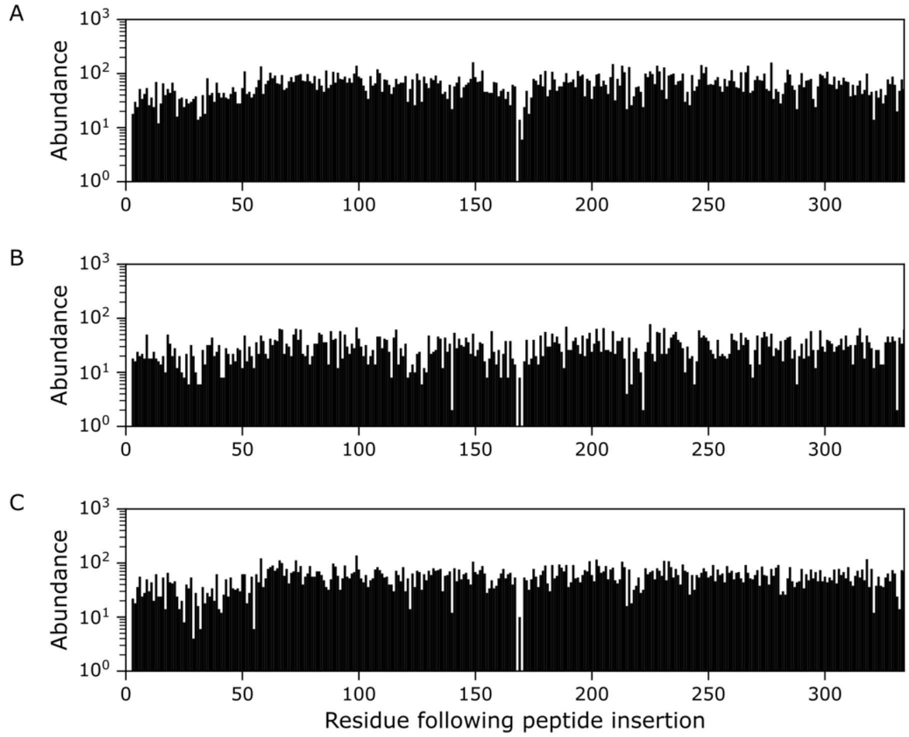


**Figure S3. Non-selected library sequence abundances in three experiments.** Raw sequencing data was filtered for reads having insertions within the MtrA sequence. The abundance of reads for each insertion variant is provided for: (**A**) replicate 1 (n = 21420 filtered reads), (**B**) replicate 2 (n = 9934), and (**C**) replicate 3 (n = 18538).

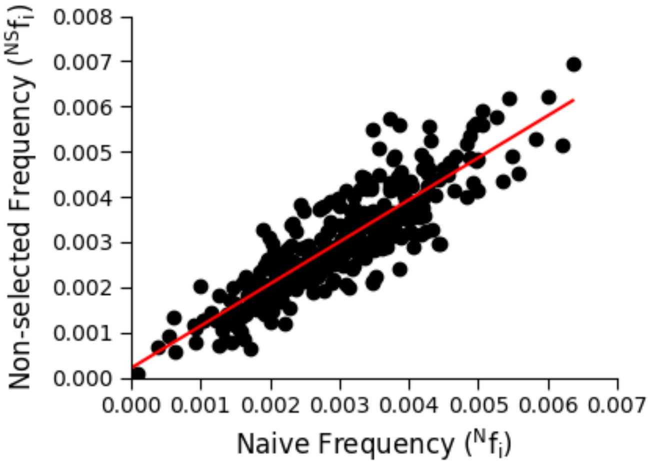


**Figure S4. A comparison of non-selected and naïve frequencies for each insertion variant.** A fit of the data using linear regression (red) yields a slope of 0.93.

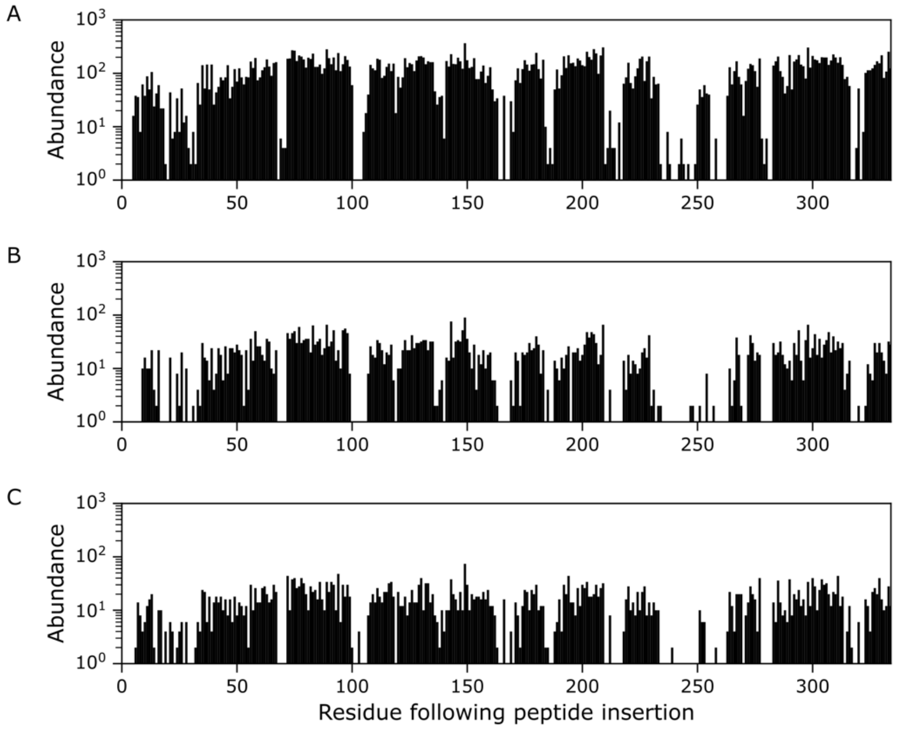


**Figure S5. Selected library sequence abundances from three experiments.** Raw sequencing data was filtered for reads with insertions within the MtrA sequence. The abundance of reads for each insertion variant is listed for: (**A**) replicate 1 (n = 32718 filtered reads), (**B**) replicate 2 (n = 5364), and (**C**) replicate 3 (n = 4452).

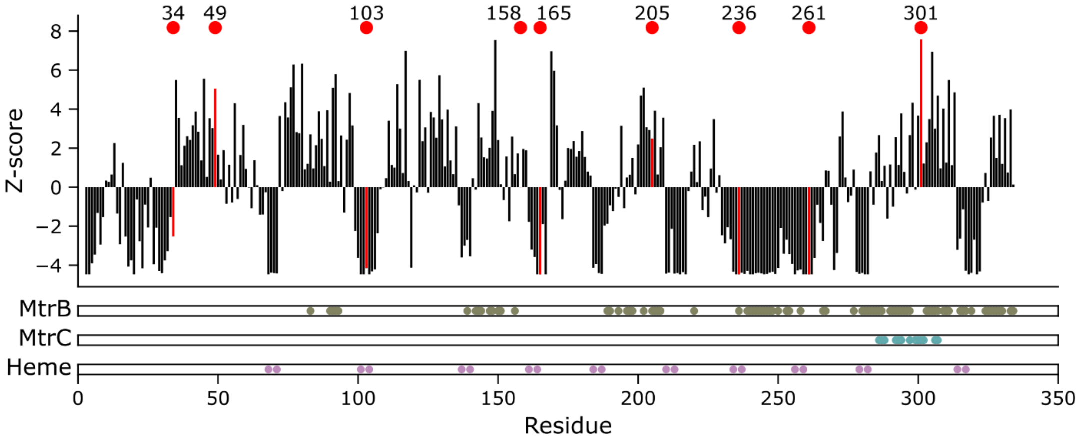


**Figure S6. Enrichment values calculated for each mutant.** The MtrA residues that make contacts with MtrB (dark green) and MtrC (teal) are colored, as well as the residues that ligate hemes (pink). Insertion variants that were characterized are highlighted in red and labeled with the residue index following their insertion.

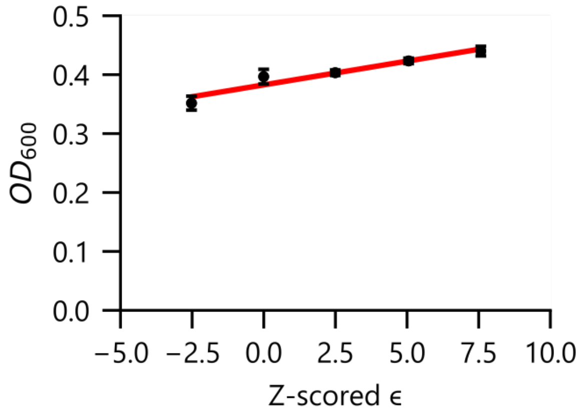


**Figure S7. Growth complementation of *So*-Δ*mtrA* by vectors expressing insertion variants with different z scores.** Cells were grown under anaerobic conditions with ferric citrate as the sole electron acceptor at 30°C. Optical densities measured after 24 hours are compared with z scores observed in the mutational profile and fit using linear regression ($r^2 = 0.93$). Error bars represent ±1σ from three experiments.

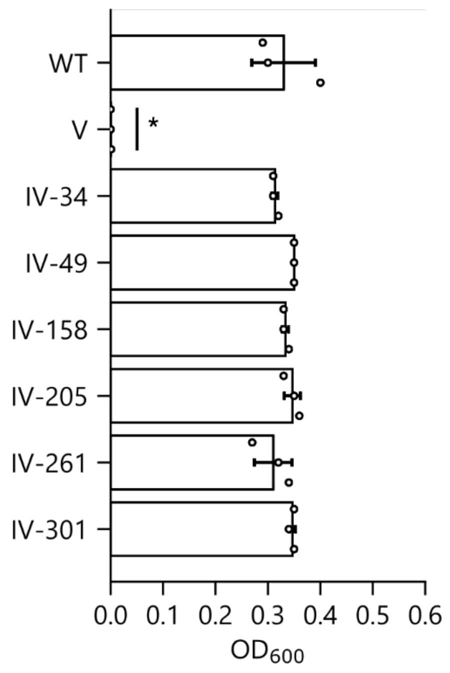


**Figure S8. Growth complementation of *So*-∆*mtrA* by MtrA insertion variants following a 72 hour incubation.** Cells were transformed with an empty vector (V), a plasmid that expresses wildtype (WT) MtrA, and plasmids that express different MtrA insertion variants. Growth was performed under anaerobic conditions with ferric citrate as the sole electron acceptor. The optical density of cultures was read at 72 hours. Error bars represent ±1 standard deviation from three experiments.

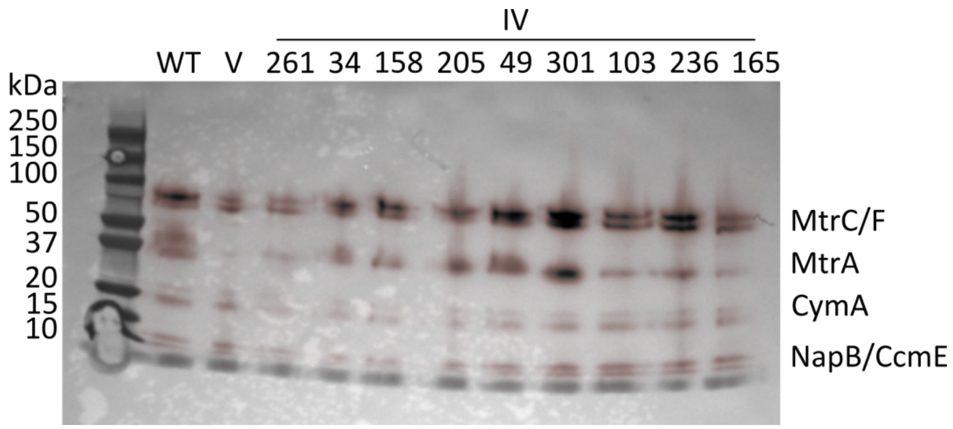


**Figure S9. Effect of mutations on cytochrome maturation evaluated using enhanced chemiluminescence.** *So*-Δ*mtrA* transformed with empty vector and plasmids expressing MtrA and different insertion variants were grown overnight in LB containing kanamycin (50 µg/mL), pelleted (4000 x g) for 10 minutes at room temperature, and washed with M9 medium. Total protein from each sample was separated using an agarose gel and stained for cytochromes. The image represents an overlay of the chemiluminescent (red) and visible (black) images. Bands corresponding to cytochromes in *S. oneidensis* and molecular weight markers are labelled.

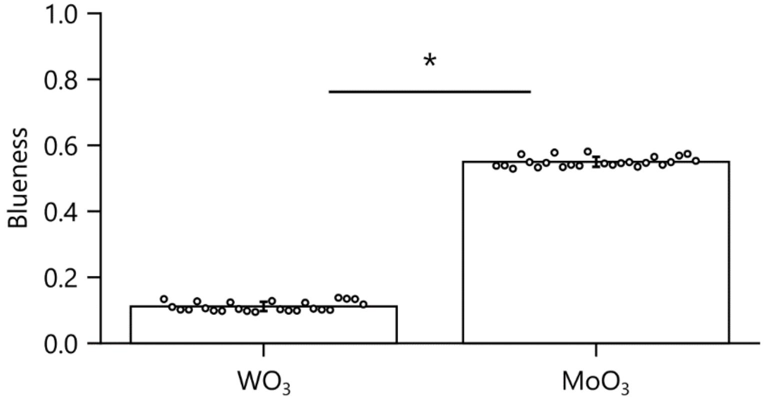


**Figure S10. A comparison of the reduction sensitivity of $WO_3$ and $MoO_3$ nanoparticles.** Saturated *Shewanella* cultures were incubated in an anaerobic chamber with 10 mg/mL of each terminal electron acceptor and then quantified for blue color intensity (n=24). The bar and asterisk denote a significant difference between the signals observed with $WO_3$ and $MoO_3$ materials after a two-tailed t-test (p value < $10^{-50}$). Error bars represent ±1 standard deviation.

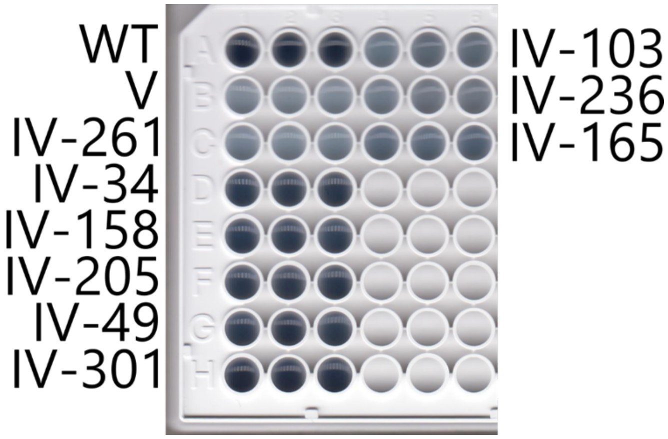


**Figure S11. Image of 96-well plate following blueness reduction measurements.** *So*-Δ*mtrA* transformed with an empty vector (V), a plasmid expressing wildtype MtrA (WT), and vectors expressing the different insertion variants were cultured overnight were incubated in minimal M9 media containing 10 mg/L molybdenum oxide ($MoO_3$) nanoparticles under an anaerobic atmosphere for 15 minutes in a 96-well plate prior to imaging. Three biological replicates were performed with each sample.

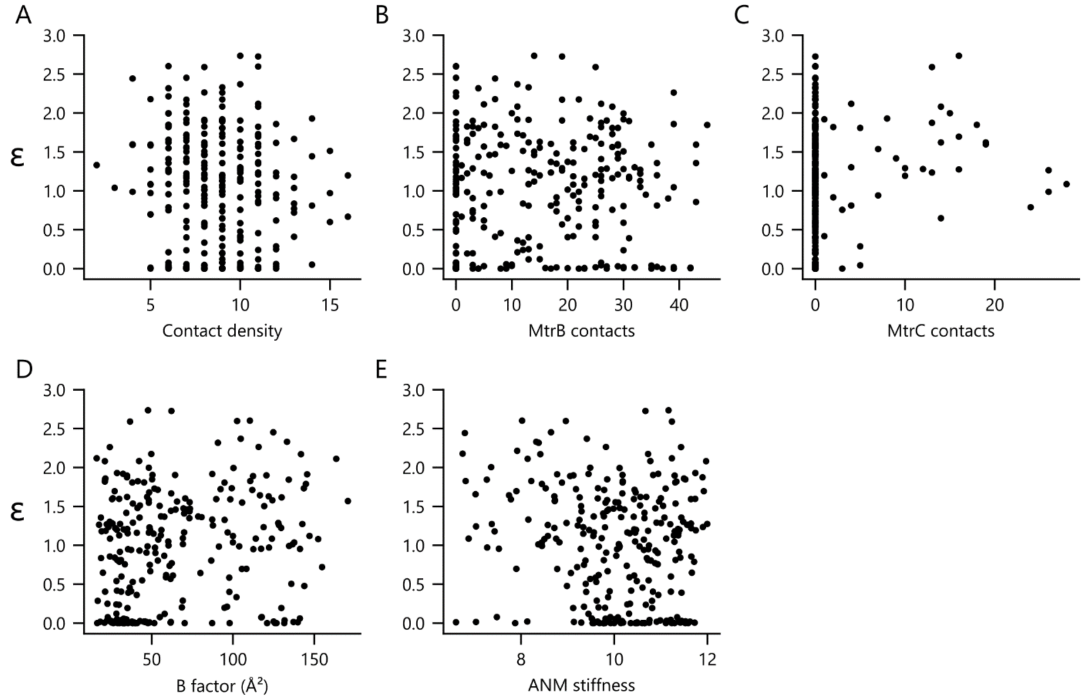


**Figure S12. A comparison of the insertional tolerance and structural features in MtrA.** Using the *S. baltica* MtrA as a model (PDB: 6r2q), several structural features were computed. Spearman correlations were calculated for comparisons of z scores with: (**A**) short-range intra-MtrA contacts, defined as ≤8 Å ($r_s$ = -0.087; p = 0.153), (**B**) contacts with MtrB, defined as ≤14 Å ($r_s$ = 0.070 p = 0.254, (**C**) contacts with MtrC, defined as ≤14 Å ($r_s$ = 0.193; p = 0.001), (**D**) averaged B-factor ($r_s$ = 0.187; p = 0.002), and (**E**) stiffness from a coarse-grain anisotropic network model ($r_s$ = -0.113; p = 0.063).

**Table S1. Assay vectors used.** For each vector, the name, antibiotic marker, origin, proteins expressed, and Addgene IDs are noted.

**Supplementary Table 1. Plasmids.** Vectors are listed with antibiotic markers, plasmid origin, and genes expressed.

| Plasmid Name | Addgene ID | Description |
|---|---|---|
| pJA018 | 179562 | $Kan^R$, RSF1010 vector with aTc inducible *S. oneidensis* MtrA |
| pIC004 | 179563 | $Kan^R$, RSF1010 empty vector |
| pIC013 | 179564 | $Kan^R$, RSF1010 vector with aTc inducible *S. oneidensis* MtrA inserted with SGRPGSLS at P261 (IV-261) |
| pIC014 | 179565 | $Kan^R$, RSF1010 vector with aTc inducible *S. oneidensis* MtrA inserted with SGRPGSLS at A34 (IV-34) |
| pIC015 | 179566 | $Kan^R$, RSF1010 vector with aTc inducible *S. oneidensis* MtrA inserted with SGRPGSLS at V158 (IV-158) |
| pIC016 | 179567 | $Kan^R$, RSF1010 vector with aTc inducible *S. oneidensis* MtrA inserted with SGRPGSLS at A205 (IV-205) |
| pIC017 | 179568 | $Kan^R$, RSF1010 vector with aTc inducible *S. oneidensis* MtrA inserted with SGRPGSLS at T49 (IV-49) |
| pIC018 | 179569 | $Kan^R$, RSF1010 vector with aTc inducible *S. oneidensis* MtrA inserted with SGRPGSLS at N301 (IV-301) |
| pIC032 | 179570 | $Kan^R$, RSF1010 vector with aTc inducible *S. oneidensis* MtrA inserted with SGRPGSLS at C103 (IV-103) |
| pIC033 | 179571 | $Kan^R$, RSF1010 vector with aTc inducible *S. oneidensis* MtrA inserted with SGRPGSLS at C236 (IV-236) |
| pIC036 | 179572 | $Kan^R$, RSF1010 vector with aTc inducible *S. oneidensis* MtrA inserted with SGRPGSLS at Q165 (IV-165) |
